# Model-driven optimal experimental design for calibrating cardiac electrophysiology models

**DOI:** 10.1101/2022.11.01.514669

**Authors:** Chon Lok Lei, Michael Clerx, David J. Gavaghan, Gary R. Mirams

## Abstract

Models of the cardiomyocyte action potential (AP) have contributed immensely to the understanding of heart function, pathophysiology, and the origin of heart rhythm disturbances. However, AP models are nonlinear, complex, and can contain more than a hundred differential equations, making them difficult to parameterise. Therefore, cellular cardiac models have been limited to describing ‘average cell’ dynamics, when cell-specific models would be ideal to uncover inter-cell variability but are too experimentally challenging to be achieved. Here, we focus on automatically designing experimental protocols that allow us to better identify cell-specific maximum conductance values for each major current type—optimal experimental designs—for both voltage-clamp and current-clamp experiments. We show that optimal designs are able to perform better than many of the existing experiment designs in the literature in terms of identifying model parameters and hence model predictive power. For cardiac cellular electrophysiology, this approach will allow researchers to define their hypothesis of the dynamics of the system and automatically design experimental protocols that will result in theoretically optimal designs.

## 1 Introduction

Cardiac cellular electrophysiology is the study of how cells manipulate ionic concentrations and membrane voltage to initiate and synchronise heartbeats. Having demonstrated their usefulness in fundamental research, quantitative models of the action potential (AP) are increasingly being used in safety-critical applications (Mirams et al., 2012; Niederer et al., 2018) such as optimising patient treatments (Corral-Acero et al., 2020; Niederer et al., 2020) or assessing drug safety (Li et al., 2019). These models contain several parameters, which may represent chemical properties such as binding rates, but also patient and even cell-specific properties such as ion current conductances (which are determined by gene expression levels). In many cases such parameters are not measured directly, but *inferred* indirectly from experimental recordings of membrane potential or transmembrane current. To create accurate models, we need to ensure the experiments used in their creation contain sufficient information on all parameters we need to infer. However, the experiments used in this process have not usually been designed with model calibration in mind, and as a result some parameters may be poorly constrained by the available data (Clerx et al., 2019a; Whittaker et al., 2020). Optimal experimental design (OED) is a method of designing experiments which uses an objective approach to elicit data containing “optimal information” for model calibration (Lindley et al., 1956; Kiefer, 1959; Smucker et al., 2018). It is extremely popular in physical sciences such as geosciences (Ushijima and Yeh, 2013; Seidler et al., 2016; Ushijima et al., 2021), mechanical engineering (Wickham et al., 1995; Gupta and Dhingra, 2013), chemical engineering (Rodriguez-Fernandez et al., 2007; Gottu Mukkula and Paulen, 2017; Schenkendorf et al., 2018), etc. In this study, we apply the techniques of OED to cardiac cellular electrophysiology to expedite the development of AP models suitable for both basic research and safety-critical applications.

The electrophysiological properties of many types of isolated cells can be studied and examined via patch clamp experiments (Hamill et al., 1981; Sakmann and Neher, 1995) in which the experimenter either controls (‘clamps’) the cell’s membrane potential and measures the resulting transmembrane current (voltage clamp mode), or controls the current while measuring potential (current clamp mode). In either mode, a pre-determined waveform (i.e. a sequence of voltages or currents) is applied, which we shall refer to as the experimental *protocol*. OED is a methodology of designing protocols based on the intuition that a model parameter can only be estimated from a measured model output (e.g. a current or voltage trace) if that output is strongly sensitive to the parameter value during the experiment. In short, it produces protocols that *maximise* (in some exact sense to be chosen by the experimenter) the model output’s sensitivity to all parameters of interest.

We apply an OED strategy to design experimental protocols that allow us to better identify cellspecific maximum conductance values for the major current types in mathematical cardiac myocyte models. Using simulation results from a controllable, understandable cellular system, we show that OED can successfully be applied to both voltage-clamp and current-clamp experiments to obtain cellspecific models. Moreover, the results demonstrate how OED protocols perform better than many of the existing experiments in the literature, in terms of reducing uncertainty in model parameters and its theoretical predictive power. This study provides a road map for researchers in cardiac cellular electrophysiology to automatically design theoretically-optimal experimental protocols.

## 2 Results

In this section we briefly introduce our OED approach (a full exposition is given in Methods) before presenting 12 new protocols derived for voltage clamp and 6 new protocols for current-clamp experiments. Various ways to evaluate the ‘quality’ of these optimised protocols are explored and we discuss some of the properties that we would like or expect them to have. To check the designs, we evaluate them using both theoretical optimality measures and practical tests in which we infer parameters from simulated data. Finally, we compare the OED performance against designs found in the literature.

### 2.1 Optimal experimental designs for patch clamp experiments

OED provides a framework for obtaining high-quality, statistically grounded designs ***u*** that are optimal with respect to a statistical criterion Φ. It is based on the assumption that we can parameterise the designs ***u*** with a *control* or *design variable vector* ***d*** such that it can be optimised with the criterion Φ that depends on the choice of a model *M*. A schematic overview of the method is given in Figure 1.

**Figure 1:**
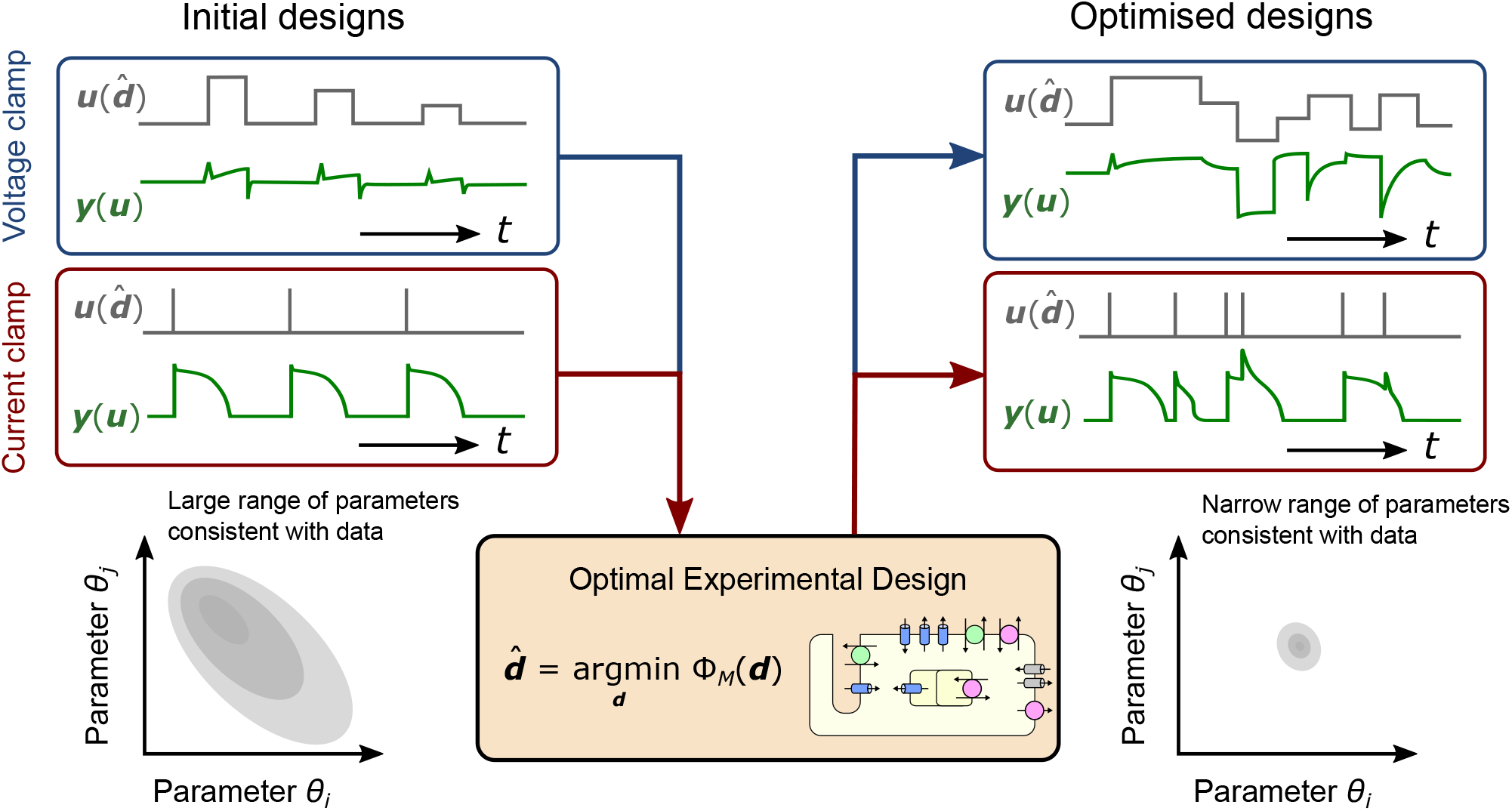
A schematic of optimal experimental design for patch-clamp experiments. Initial designs (left, ***u***(***d***)) of the voltage-clamp and current-clamp experiments, which can commonly be found in the literature or perhaps are ‘intuitive’ choices of the design inputs (***d***), elicit uninformative data/observables ***y***(***u***) for calibrating complex mathematical models, leading to a wide posterior for the model parameters ***θ*** (shown for two parameters but the principle applies for higher numbers of parameters). Optimal experimental design with the mathematical model *M* under a certain statistical criterion Φ_*M*_ derive optimised designs (right, 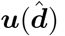) with non-intuitive choices of design inputs 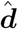 which elicit informative data for calibration, and lead to narrower parameter posteriors.

Several options have been proposed for Φ, usually relating to the remaining uncertainty in model parameter values ***θ*** after calibration, or to other statistically relevant properties of the design. The criteria used in this study are known as the A-criterion, the D-criterion, and the E*-criterion (see § 4.3). They can be interpreted as properties (the shape and size) of the *confidence hyper-ellipsoid* for the calibrated model parameters ***θ*** (Vanrolleghem and Dochain, 1998; Banga and Balsa-Canto, 2008): the A-optimal design can be interpreted as minimising the average variance of the confidence ellipsoid; the D-optimal design minimises the volume of the confidence ellipsoid; and the E*-optimal design minimises the ratio of the length of the largest to the smallest axis, making the ellipsoid as spherical as possible. The confidence ellipsoid were approximated using the Fisher information matrix based on the sensitivity of the model *M* (see § 4.3). Two types of sensitivity were employed: local sensitivity analysis (LSA) calculates the sensitivity of model outputs for a *given* set of parameters ***θ*** (using the default model parameters from the literature), and hence its name ‘local’ (i.e. optimal in a region of parameter space which is close, or local, to the assumed parameters); on the other hand global sensitivity analysis (GSA) samples parameters across a predefined parameter space (here we used a hypercube where each model parameter *θ*_*i*_ can be *e*−2 to *e*−2 times its original value).

Since the statistical criterion is evaluated based on the choice of a model, Φ_*M*_, we first considered using the adult human ventricular (epicardial) AP model by O’Hara et al. (2011) to calculate the criterion. We further considered using the *average* of the statistical criteria 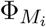 calculated using multiple models (*M*_*i*_ = {*M*_1_, *M*_2_, …}), with the aim of obtaining a design that is optimal for a range of biological assumptions instead of focusing on one particular model, termed the ‘averaged model criterion’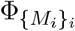. Here we used five adult ventricular (epicardial) AP models to represent an averaged behaviour: ten Tusscher et al. (2004), Fink et al. (2008), O’Hara et al. (2011), Dutta et al. (2017) and Tomek et al. (2019). This gives us two types of model for calculating the criterion: single model criterion and averaged model criterion.

Two types of experimental protocols (***u***) are optimised: voltage clamp and current clamp (see § 4.2). We defined voltage clamp protocols as sequences of steps with a magnitude and duration that could be varied by OED as the design variables ***d***. Our current clamp protocols consisted of short bursts of current injection with a fixed magnitude and duration, here the interval between burst was used as design variable (see Figure 1). OED can also work within a variety of constraints, such as restrictions in duration of the experiment or applied voltage ranges, to ensure the optimal designs are practically feasible.

Experiment designs ***u***(***d***) were optimised based on a design measure which consists of a choice of the optimal design criterion and a choice of model used to perform the design, using a global optimisation algorithm. For voltage clamp experiments, we used six design criteria (LSA A-, D-, E*-designs, and GSA A-, D-, E*-designs) and two types of model, giving us in total 12 optimal design strategies (see § 4.3). The resulting optimised protocols using different strategies are shown in Figure 2 (Protocols A–L), with the corresponding current (***y*** = *I*_*obs*_) simulated using the O’Hara et al. (2011) model (the corresponding eigenvalue spectra are shown in Supplementary Figures). For current clamp experiments, we used three design criteria (the LSA A-, D-, E*-designs) and two types of model, giving us in total 6 optimal design strategies. The resulting optimised protocols using different strategies are shown in Figure 3 (Protocols O–T) with the corresponding membrane voltage (***y*** = *V*_*m*_) simulated using the O’Hara et al. (2011) model.

**Figure 2:**
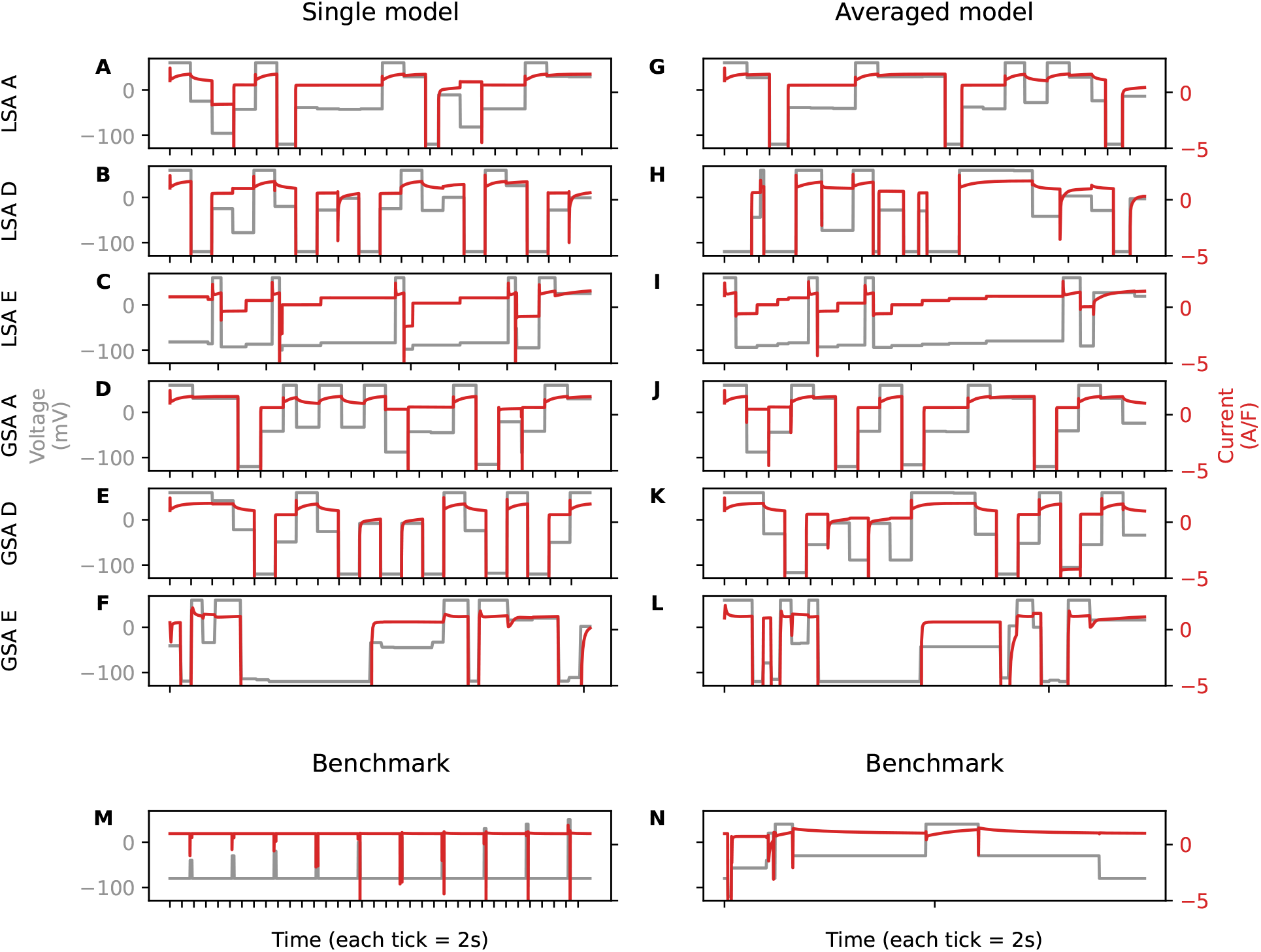
**(A–L)** The optimised protocols (shown in grey, values on left axis) using different criteria and the corresponding *I*_*obs*_ (shown in red, values on right axis) simulated using the O’Hara et al. (2011) model. Rows are protocols optimised with different optimal design criteria: the LSA A-, D-, E*-designs, and the GSA A-, D-, E*-designs. Columns are protocols optimised based on different types of models: the single model, and averaged model. **(M, N)** The benchmark protocols from Lei et al. (2017) and from Groenendaal et al. (2015).

**Figure 3:**
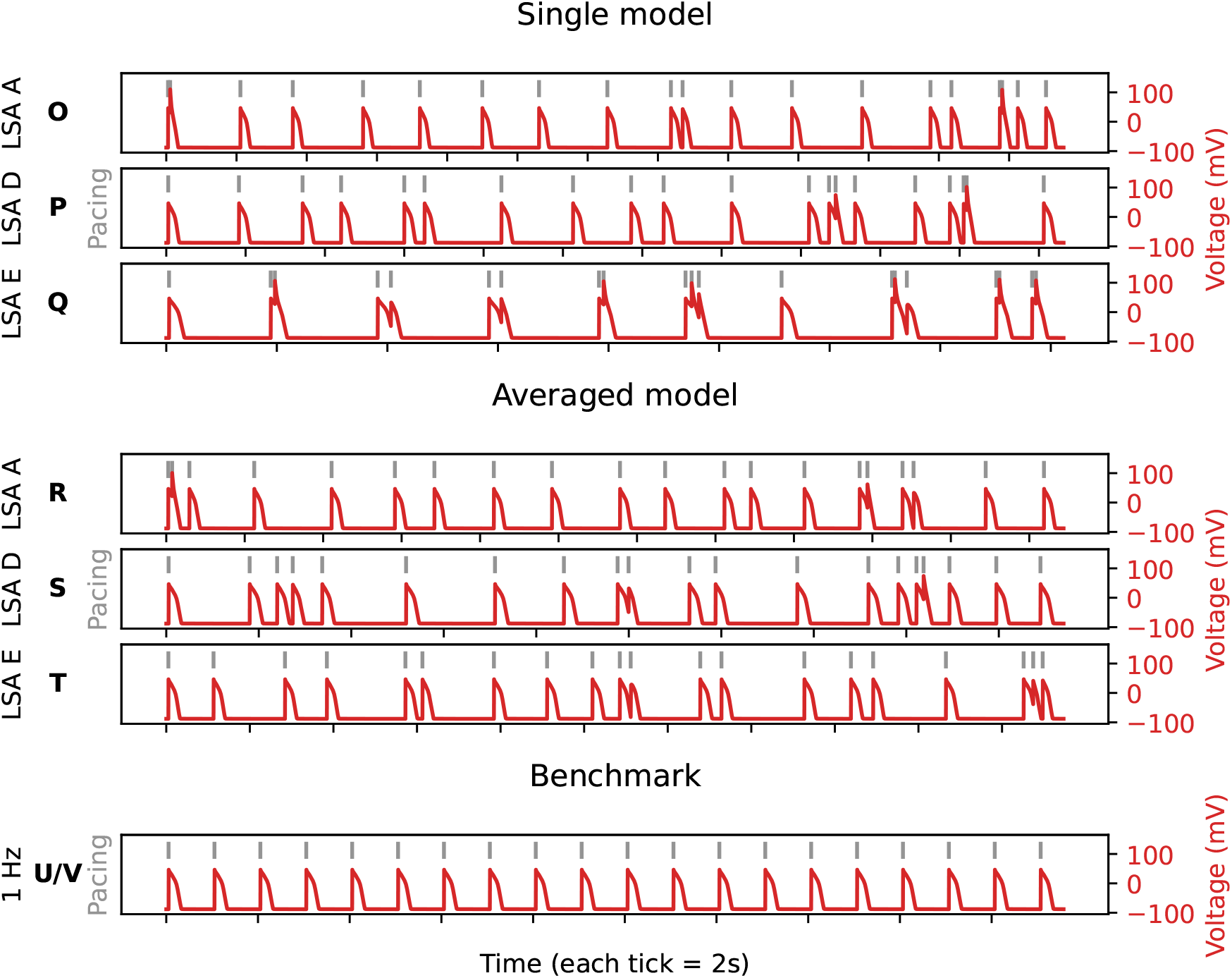
The optimised protocols (shown in grey, the positions at which stimuli were applied) using different criteria and the corresponding membrane voltage *V*_*m*_ (shown in red, values on right axis) simulated using the O’Hara et al. (2011) model. Rows are protocols optimised with different optimal design criteria: the LSA A-, D-, E*-designs. Protocols were optimised based on **(O–Q)** single model, and **(R–T)** averaged model. **(U/V)** The benchmark protocols (1 Hz pacing).

Each of the protocols appears to elicit rich and varied dynamics. Since these optimal protocols were designed to maximise the information for all model parameters for inference, we evaluate the performance of these optimised protocols and compare them against some protocols from the literature.

### 2.2 Do OEDs help us find model parameters?

Taking a practical approach, we generated synthetic (simulated) data under these optimised protocols 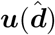, then asked the following questions: (i) can we identify the model parameters ***θ*** correctly using these synthetic data? (ii) how does the error, if any, and uncertainty in the parameters vary between these optimised protocols?

We generated synthetic data using the O’Hara et al. (2011) model with i.i.d. Gaussian synthetic noise 𝒩 (0, *σ*^2^) with zero mean and standard deviation *σ* = 0.15 A F^−1^ for voltage clamp and *σ* = 1 mV for current clamp. Then the same model was used to fit the synthetic data, to test whether we can identify the maximum conductance parameters ***θ*** when we have a good estimate of the kinetics. Posterior distributions of the parameters were estimated using a Markov chain Monte Carlo (MCMC) based sampling scheme (Jasra et al., 2007), where the final chains converged with 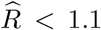 for all parameters (Gelman et al., 2013).

In terms of reducing the uncertainty and error in the inferred parameters, the two most successful optimal designs 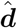 for voltage-clamp experiments were the LSA D-design and GSA A-design for the O’Hara et al. (2011) model (Protocols B and D), as shown in Figure 4A. Each chain returned a posterior distribution over the parameters, since all chains converge and they are all very similar to each other, we use all three chains to compute the marginal parameter distributions (per parameter). The mean (over all the parameters) root-mean-square error (RMSE) of these distributions to the true values is shown in Figure 4B for all of the voltage-clamp protocols.

**Figure 4:**
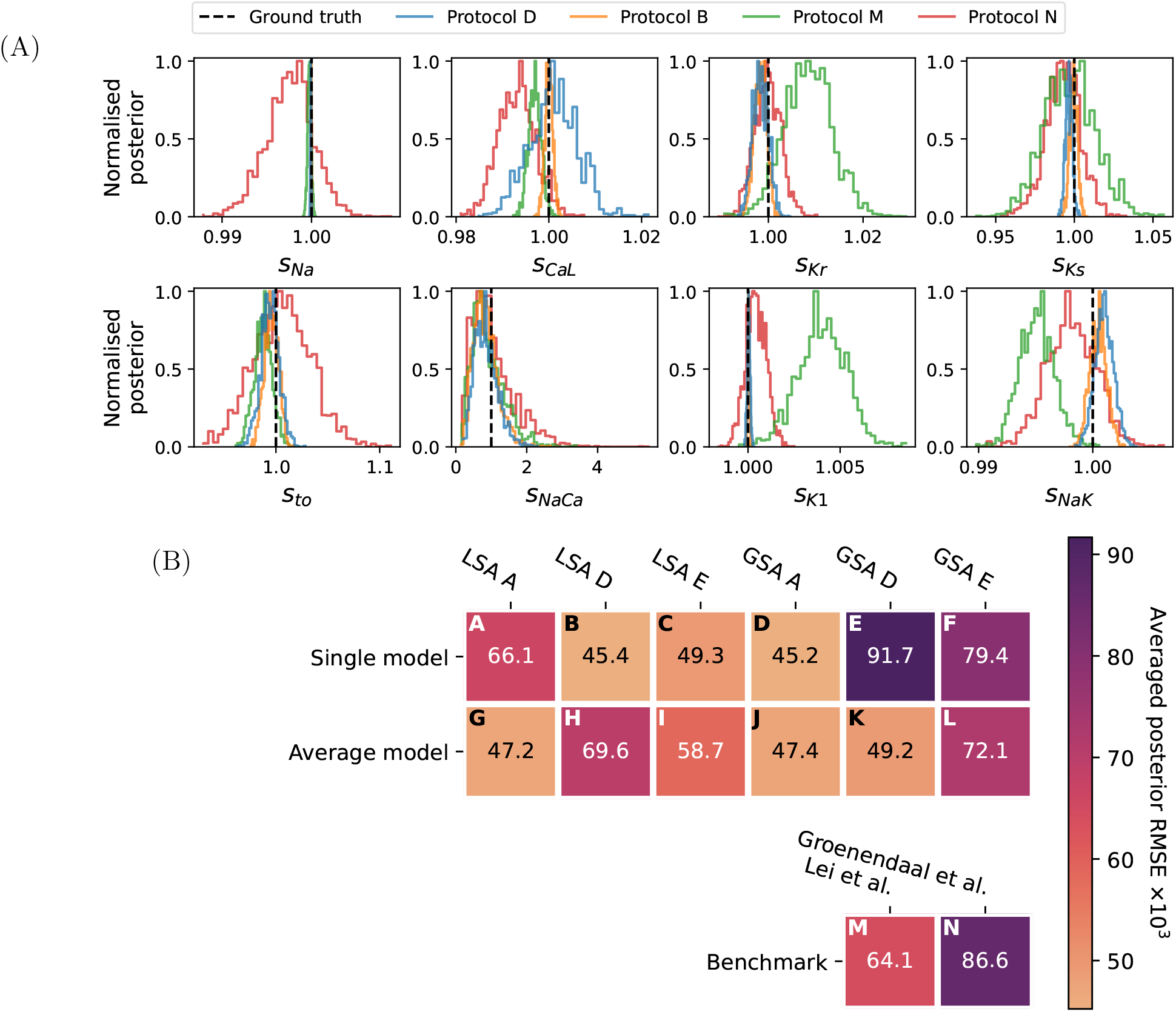
The posterior distribution of the parameters for the synthetic study with voltage clamp, generated with the O’Hara et al. (2011) model and fitted with the same model. **(A)** The marginal posterior for the best two protocols: LSA D- and GSA A-designs for the O’Hara et al. (2011) model (Protocols B and D), according to the mean RMSE of the marginal posterior to the true values, and for the two benchmark protocols. **(B)** The mean RMSE of the marginal posterior for all of the optimised protocols, as well as the benchmark protocols.

For current-clamp experiments, all the OED results had a similar posterior width, with the most successful optimal design to be the LSA A-design for the single model (Protocol O) as shown in Figure 5A. The mean (over all the parameters) RMSE of these distributions is shown in Figure 5B for all of the current-clamp protocols.

**Figure 5:**
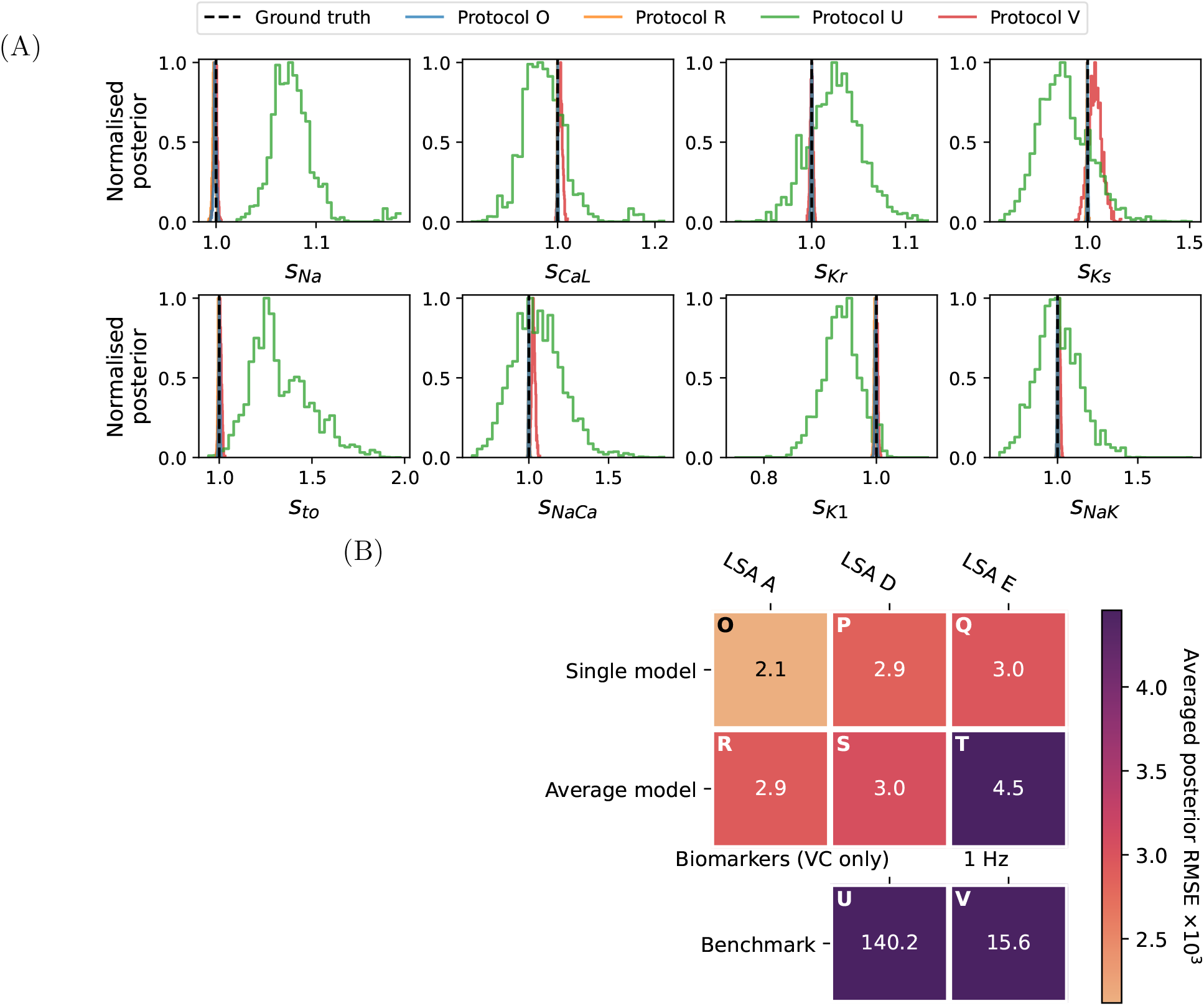
The posterior distribution of the parameters for the synthetic study with current clamp, generated with the O’Hara et al. (2011) model and fitted with the same model. **(A)** The marginal posterior for the best two protocols: LSA A-design (Protocols O and R), according to the mean RMSE of the marginal posterior to the true values, and for the two benchmark protocols. Note that y-axes are rescaled to unity for visualisation purpose. Magnification of this figure around true value is provided in Supplementary Figures. **(B)** The mean RMSE of the marginal posterior for all of the optimised protocols, as well as the benchmark protocols.

### 2.3 Cross-measure assessments of OEDs

We may also expect the best protocol from those OEDs would be a robust good ‘all-round’ protocol. Each of the OED is by definition optimal under a given design measure, whilst a robust good all-round design would not be the worst under another design measure and, on average, should be good under all measures.

The performance of all the optimised protocols under a particular experimental mode were compared across all the design measures; a design measure is defined by the choices of the model and the design criterion. We used the ten Tusscher et al. (2004) model, the Fink et al. (2008) model, the O’Hara et al. (2011) model, the Dutta et al. (2017) model, the Tomek et al. (2019) model, and the averaged model to assess the OED performance; the O’Hara et al. (2011) model and the averaged model were used to optimise the protocols, and the remaining five were included to assess how these optimised protocols perform should data really arise from cells that are better described by these other models. We used the same criteria to assess the OEDs, and their combinations form what we refer to as the cross-measure matrix. Each design measure entry of the cross-measure matrix was normalised to 0–100 % with the best and the worst values seen across all the OED protocols such that it can be compared with other design measures.

Examples of cross-measure assessments for the OEDs are shown in Figure 6A for voltage clamp and Figure 7A for current clamp. Figure 6A shows our best voltage clamp protocol design, the OED with the ‘averaged model score’ under GSA D-criterion (Protocol K), which performs well under all measures and model currents. Figure 7A shows our best current clamp protocol design, the OED with the ‘averaged model score’ under LSA A-criterion (Protocol R), which performs well under all measures and model currents. The ‘best’ protocols in this assessment were decided based simply on the mean of all entries in the cross-measure matrix. The averaged score for each optimised protocol is shown in Figure 6C for voltage clamp and Figure 7C for current clamp.

**Figure 6:**
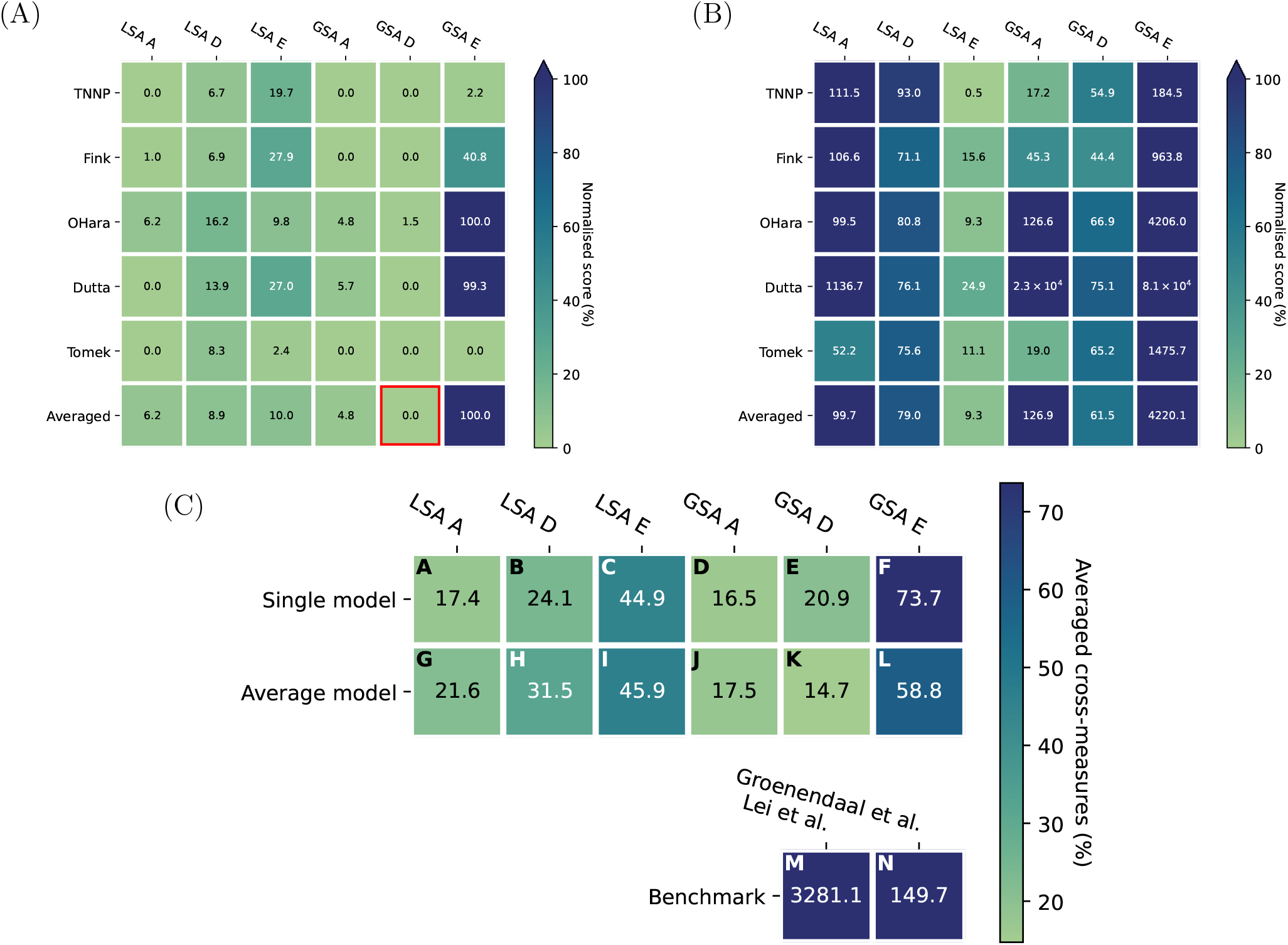
Cross-measure evaluations for the optimised voltage-clamp protocols, compared against the benchmark protocols. **(A)** The normalised cross-measure matrix for the best protocol, Protocol K, optimised using the GSA D-criterion and the averaged model (indicated with the red box). **(B)** The matrix for the worst protocol, Protocol M, from Lei et al. (2017). **(C)** The mean of the normalised cross-measure evaluations for all of the optimised protocols, as well as the benchmark protocols.

**Figure 7:**
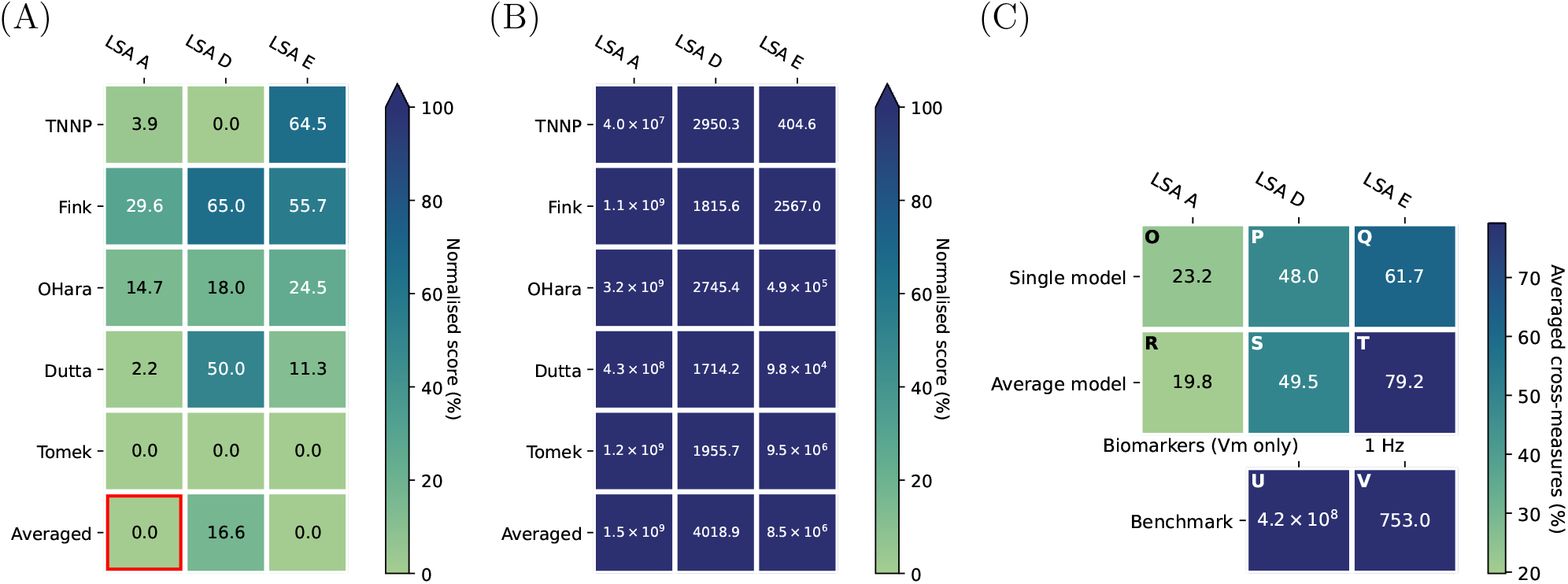
Cross-measure evaluations for the optimised current-clamp protocols, compared against the benchmark protocols. **(A)** The normalised cross-measure matrix for the best protocol, Protocol R, optimised using the LSA A-criterion and the averaged model (indicated with the red box). **(B)** The matrix for the worst protocol, Protocol U, biomarker estimation with 1 Hz pacing. **(C)** The mean of the normalised cross-measure evaluations for all of the optimised protocols, as well as the benchmark protocols.

Figure 6C and Figure 7C show that the protocols based on multiple models (bottom row) are similar to those based on only the O’Hara et al. (2011) model (top row), although the best protocols for each experimental mode is based on multiple models. This may be expected as the protocols based on only one model may perform relatively badly under other models. We also noticed that the protocols based on the LSA E criterion are worse than the other LSA criteria. Finally, in Figure 6C, the protocols based on the GSA criteria are better than (or similar to) their LSA counterparts, apart from the E*-design using the O’Hara et al. (2011) model.

### 2.4 OEDs outperform literature experimental designs

Finally, we assessed the OED results by comparing their performance against some of the designs of experiments in the literature—a benchmark comparison. For voltage-clamp experiments, we used (1) a simple step protocol from Lei et al. (2017) and (2) an expertly-designed protocol from Groenendaal et al. (2015) to benchmark the performance. For current-clamp experiments, we used (1) a simple 1 Hz constant pacing protocol; or (2) derived summary statistics (biomarkers) based on 1 Hz pacing, which are commonly used in the literature (Bartolucci et al., 2020; Britton et al., 2013; Jæger et al., 2019, 2020, 2021; Paci et al., 2017, 2018; Tveito et al., 2018).

For the voltage-clamp protocols, the benchmark protocol from Lei et al. (2017) (Protocol M) shown in Figure 6B performed badly for most of the design measures, which may be expected as it is not designed using any of these measures. Similarly, the protocol from Groenendaal et al. (2015) scored badly compared to OED protocols, although its score was, on average, just twice that of the worst OED protocol (Protocol F) which is impressive given its manual design. Figure 4 also shows that the benchmark protocols resulted in considerably wider posterior distributions than most of the optimized protocols. Often the number of data samples (or data points) in the i.i.d. Gaussian likelihood that we use is a major factor determining the width of the posterior. Here even for the protocol from Lei et al. (2017) (Protocol M) which is much longer in duration (hence more data samples) than any of the optimised protocols, the width of its posterior is still bigger than most of the optimised protocols. Interestingly, even though the protocol by Groenendaal et al. (2015) (Protocol N) was designed specifically to tease out different dynamics of the ionic currents, the width of the posterior is similar to that of the protocol from Lei et al. (2017). Overall, this shows that the OED methods were successful in reducing the parameter uncertainty for voltage clamp experiments.

For current clamp experiments, the biomarkers that we used were action potential duration (APD) at 10 % repolarisation (APD10), APD20, APD30, APD40, APD50, APD60, APD70, APD80, APD90, Triangularisation (Tri, defined as APD90 − APD40), plateau duration, area under the curve of voltage trace (at APD30), maximum upstroke velocity, resting potential, maximum voltage, dome peak, and action potential amplitude (APA) which were used in Bartolucci et al. (2020); Britton et al. (2013); Jæger et al. (2019, 2020, 2021); Paci et al. (2017, 2018); Tveito et al. (2018); to perform practical evaluations for biomarkers, we used a percentage difference for calculating the likelihood as it was done similarly in e.g. Tveito et al. (2018); Jæger et al. (2019). We first note that the same was observed for the 1 Hz pacing protocol: not only is the cross evaluation matrix consistently worse than the OED protocols (Figure 7B), but the posterior obtained in the practical assessment is also wider than all of the OED protocols (Figure 5). Moreover, the results from using the biomarkers (Figure 5A) were not only more uncertain than the other approaches but also biased in some of the parameter estimates, e.g. *I*_*Na*_ and *I*_*to*_. The results suggest that the OED protocols were much better in estimating the model parameters than those in the literature.

## 3 Discussion

We have applied OED to various AP models to design voltage-clamp and current-clamp protocols for fitting maximum conductances. This is, to our knowledge, the first time such an approach has been taken in cardiac cellular electrophysiology. Some studies have looked into designing experiments for model calibration from *expertly* designed protocols, ranging from calibrating models of individual ionic current dynamics (Fink and Noble, 2009; Zhou et al., 2009; Beattie et al., 2018; Lei et al., 2019b,a) to whole-cell models of APs of cardiomyocytes (Groenendaal et al., 2015); similarly in neuroscience (Hobbs and Hooper, 2008; Tomaiuolo et al., 2012) too. However, these protocols are not truly ‘optimal’ under a certain statistical or objective measure; they are designed through (subjective) expert views. Our (objectively) optimised protocols successfully reduced the uncertainty in the inferred parameters, compared to some of the conventional protocols.

We explored various designs adapted from the OED literature. Most of them were able to improve the uncertainty in the inferred parameters compared to the benchmark protocols. We tried to seek the best protocol amongst the OED generated protocols. However, we were not able to identify the best one, as each of them perform slightly differently under different situations. For example Protocol K was the best in the cross-measures comparison, whilst Protocol D was better in reducing parameter uncertainty and error. We found that the protocols based on averaged model were consistently (slightly) better than the others, which may be a better way for designing patch-clamp protocols, which implicitly makes some allowance for differences in ion current kinetics.

Fitting conductances to whole-cell recordings while using ion channel kinetics models taken (or slightly adapted) from the literature, has been commonly used (Whittaker et al., 2020). Many inference approaches and biomarkers have been adopted, including multivariate regression (Sarkar and Sobie, 2010), history matching (Coveney and Clayton, 2018), “population of models” (Muszkiewicz et al., 2016), and moment-matching (Tixier et al., 2017), and all of them assumed “out of the box” or only slightly modified kinetics models when fitting maximum conductances. Similarly Johnstone et al. (2016) adapted a Bayesian approach with MCMC to infer maximum conductances using AP recordings. Groenendaal et al. (2015) proposed to use (cell-specific) voltage-clamp and current-clamp data with a genetic algorithm to find maximum conductances in an AP model. Some of these methods were included in our benchmark comparison for our approach; we have shown that optimal designs outperform these literature methods in terms of recovering parameters from simulated data.

Amongst all the methods that we compared, the biomarker approach performed the worst, even though it has been one of the most popular approaches. We further note that some of the biases in the biomarker posteriors were due to the biases introduced in estimating the biomarkers with noisy data. Such an issue is illustrated in Figure 8A, where biomarkers were estimated for 1000 noise realisations and compared against the biomarkers taken from idealised noise-free data (dashed line). For example, there are no simple ways of estimating the unbiased (i.e. neither overestimating nor underestimating) maximum voltage or the AP amplitude with noisy AP traces; if we simply define the maximum voltage as the maximum voltage value of the noisy data, as done here, then we will overestimate the actual maximum voltage most of the time. This is because we have a number of data samples (sampling time points) around the underlying maximum voltage where the noise is centred, then the noisy data will have values bigger (as well as smaller) than the mean most of the time, giving an overestimated maximum voltage value. However, if we define the maximum voltage as the averaged value over a (short) period of time around the maximum voltage value of the noisy data, then the maximum voltage will be biased differently depending on the duration of the averaging period and the shape of the AP. Therefore, depending on the data postprocessing procedure, the biomarkers, and hence model parameters inferred from these, can be biased differently.

**Figure 8:**
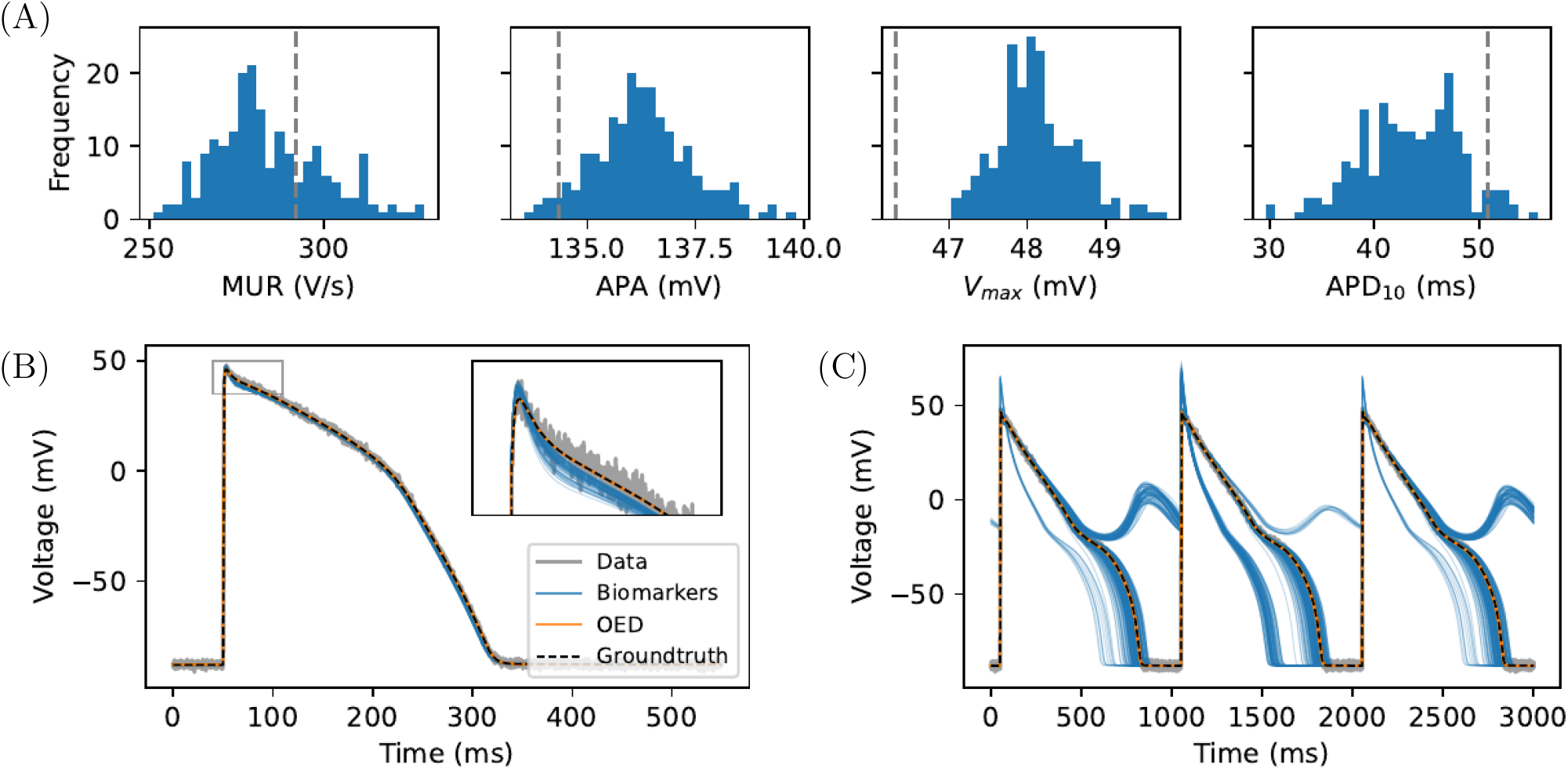
Undesired effects of using biomarkers for AP parameter inference. **(A)** Various biomarkers were estimated for 1000 noise realisations and compared against the biomarkers estimated without noise (dashed line). **(B)** Posterior predictions using the parameter posterior samples shown in Figure 5 (blue for Protocol U, biomarkers and orange for Protocol O, OED) for 1 Hz prepaced AP. **(C)** Posterior predictions for 1 Hz prepaced AP under a 90 % reduction of *I*_*Kr*_ together with a 50 % reduction of *I*_*Ks*_.

The biases introduced during biomarker extraction therefore affect posterior predictions. Figure 8B shows such posterior predictions using the parameter posteriors shown in Figure 5, where grey and black (dashed) lines show the data and the underlying ground truth, blue and orange show the predictions from the posteriors estimated using Protocol U (biomarkers) and Protocol O (LSA A with single model). The inset of Figure 8B shows a magnification around the apex of the AP, highlighting the biases of the posterior predictions with biomarkers (blue) as compared to the OED results (orange). These biases may in turn cause issues during model predictions for unseen situations. We illustrate these issues in Figure 8C which attempts to predict the AP under a 90 % reduction of *I*_*Kr*_ together with a 50 % reduction of *I*_*Ks*_. The huge variability and inaccuracy of the posterior prediction using biomarkers (blue) are consequences of both increased *parameter unidentifiability* as well as bias discussed above, as shown by comparison with the predictions with parameters inferred by fitting to the full time traces from Protocol O (orange).

Several limitations of this work suggest future studies to advance the design of experimental protocols. First, all existing OED approaches, to our knowledge, provide optimal designs only under the assumption of having a correct underlying model, meaning when there is discrepancy between the model we use and the real system, including experimental artefacts (Lei et al., 2020a, 2021), no guarantee that the designs will work better, or even work well, as discussed in (Lei et al., 2020b). The results, however, have generated experimental protocols that can readily be tested experimentally, especially since these designs can be implemented in most cellcular electrophysiology laboratories. A second limitation is that although we tested 18 OEDs, most of these are ‘classical’ design criteria that relies on certain approximation of the covariance of the estimates. In future work, we intend to determine whether more complex criteria or approaches, such as Bayesian decision theoretic approach (Huan and Marzouk, 2013; Liepe et al., 2013) and reinforcement approaches (Treloar et al., 2022), can improve the protocol design.

In summary, we have demonstrated that using mechanistic mathematical modelling and optimal experimental design can improve identification of model parameters and hence its theoretical predictive power. This work offers a methodology to automatically, objectively develop experimental protocols that can alleviate the issue of model parameter unidentifiability when collecting new experimental data.

## 4 Methods

All code and data are freely available at: https://github.com/chonlei/action-potential-experimental-design.

In the design of experiments, the results of OED are experimental designs that are optimal with respect to some statistical criteria—*optimal design measures*. Formally, we consider a (controlled) nonlinear differential equation model of the form

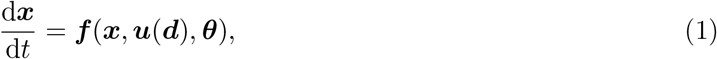

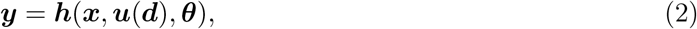

where ***x*** is the vector of model states, ***u*** is the vector of system inputs, ***θ*** is the vector of model parameters. ***f*** and ***h*** are the systems of equations, where ***f*** describes the dynamical equations, and ***h*** maps the solutions of the dynamical equations to the vector of observables ***y***, i.e. the model outputs that we compare with data. We assume that the system input ***u*** can be parameterised with ***d***, i.e. ***d*** is the vector of parameters that controls the experimental conditions. For example, if ***u*** is an applied voltage-clamp protocol, ***d*** could be the voltages and durations of its steps; if ***u*** is an applied current stimulus, ***d*** could be the durations between each stimulus, as described in details in the next sections.

The optimisation of the experimental design procedure is then defined as

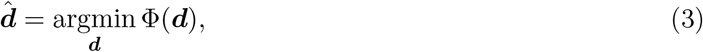

where the function Φ represents the optimal design measure to be minimised.

All optimal design measures discussed in this paper assume the model calibration process to be least-square estimation, maximum likelihood estimation, or posterior estimation; if one considers a different calibration process, such as bounded-error parameter estimation (also known as guaranteed parameter estimation) or approximate Bayesian computation (ABC), biases may be introduced, and a different set of design measures should be used (Gottu Mukkula and Paulen, 2017). All methods are implicitly conditional on the chosen set of model equations, i.e. they are only optimal for the models used during the design; this is a general limitation of model-based OED.

### 4.1 Action potential models for OED

We consider cardiac cellular electrophysiology models of APs under either a voltage-clamp or a currentclamp experiment, and we aim to optimise either the voltage-clamp or the current-clamp protocol for conductance identification. In practise, we design experiments and perform the fitting to data with a proposed model which would be our current best representation of reality.

#### Designing experiments with a single model

Here, we first use a widely-used adult ventricular (epicardial) AP model by O’Hara et al. (2011) as a ‘proposed model’. Under the voltage-clamp configuration, the observed current *I*_*obs*_ can be expressed as

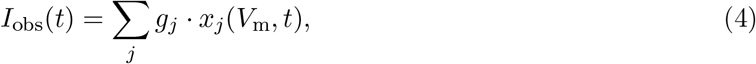

where *g*_*j*_ is the maximum conductance or permeability and *x*_*j*_(*V*) is some nonlinear function of the voltage *V*_*m*_ for the current of type *j*, and we consider the major currents only (Eq. (6)). These nonlinear functions *x*_*j*_(*V*) are the product of the ‘kinetics’ (describing the voltage-dependent opening and closing of ion channels in response to changes in membrane voltage) and the ‘driving term’ of the currents (either the Ohmic *V*_*m*_ − *E*_*S*_ term or the GHK flux equation). In this case, the observed current *I*_*obs*_ is the output defined in Eq. (2). The vector of system inputs ***u*** describes the applied voltage-clamp protocol *V* (*t*) which we assume to be the same as *V*_*m*_.

Alternatively, under the current-clamp configuration, the observed membrane voltage *V*_*m*_ follows

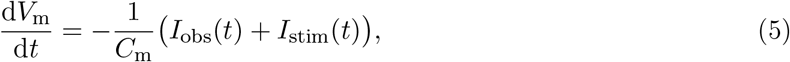

where *I*_*obs*_(*t*) is the same as Eq. (4). In this case, the membrane voltage *V*_*m*_ is the output defined in Eq. (2). The vector of system inputs ***u*** describes the applied current-clamp protocol, i.e. the stimulus current *I*_*stim*_(*t*).

We *assume* that these kinetics terms are known (and correct). We are interested in finding all *g*_*j*_, which determine the magnitude of the currents when the channels are fully open, therefore we have

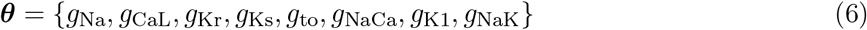

to be inferred, which are the maximum conductance or permeability of the fast sodium current, the L-type calcium current, the rapid-delayed rectifier potassium current, the slow-delayed rectifier potassium current, the transient potassium current, the sodium-calcium exchanger current, the inward rectifier potassium current, and the sodium-potassium pump current, respectively. We note that this ignores some smaller background currents in the model.

Simulations were run using Myokit (Clerx et al., 2016), with tolerance settings for the CVODE solver (Hindmarsh et al., 2005) set to abs_tol = 10^−8^ and rel_tol = 10^−10^.

#### Designing experiments with an averaged measure over models

Each OED is optimal for the model used to perform the design. However, we may want to obtain a design that is optimal for a range of biological assumptions instead of focusing on one particular model, so that the same data can be used to study multiple similar models. Therefore, we also consider using multiple models for calculating the cost function; instead of using only one model to optimise the experiment while the true data generating process could potentially be more similar to another model. That is, if we have *N*_*M*_ proposed models, then each model output *I*_*obs,M*_ (for voltage clamp) or *V*_*m,M*_ (for current clamp) is used to calculate an optimal design measure Φ_*M*_, and we optimise the experimental protocol parameters by minimising the mean of the measures:

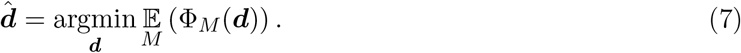

Here we use five adult ventricular (epicardial) AP models to represent an averaged behaviour: ten Tusscher et al. (2004), Fink et al. (2008), O’Hara et al. (2011), Chang et al. (2017), and Tomek et al. (2019).

### 4.2 Experiments to be optimised: Two modes of experiments

We design the experiments by varying the system input ***u***(***d***). As described in the model section, we could measure the system with two modes: voltage clamp and current clamp.

#### Designing voltage clamp experiments

The voltage-clamp protocol *V* is the system input ***u***(***d***), and we choose it to be a piecewise function defined as

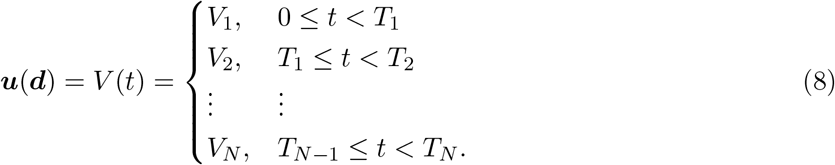

The vector of parameters that defines the voltage-clamp protocol is

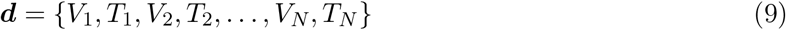

for a *N* -step protocol; equivalently we can define Δ*T*_*i*_ = *T*_*i*_ − *T*_*i*−1_ with *T*_0_ = 0, then we obtain

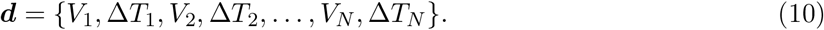

In order to ensure the resulting voltage-clamp protocol *V* is feasible to be run experimentally, we constrain the protocol to have *V*_*i*_ ∈ [−120, 60] mV and Δ*T*_*i*_ ∈ [50, 2000] ms for all *i*, which defines the protocol parameter space. We choose *N* to be 20, giving a maximum protocol duration of 40 s. We applied both the LSA and GSA designs for this experimental setting.

In theory, the voltage-clamp protocol can be replaced with any arbitrary function, e.g. a sum of sinusoidal functions (Beattie et al., 2018), as long as it can be parameterised with some vector of parameters ***d***. However, the benefit of choosing such a piecewise step function is that it can be directly applied in any patch-clamp system, including some of the high-throughput automated patch-clamp systems such as the SyncroPatch from Nanion Technologies (Lei et al., 2019b,a).

#### Designing current clamp experiments

In the current-clamp mode, the stimulus current *I*_*stim*_ is the system input ***u***(***d***). As partly inspired by Groenendaal et al. (2015), instead of *randomly* applying the stimulus current, here we *optimise* when to apply the stimulus current. We parameterised the current-clamp protocol as the following, the stimulus current with an amplitude 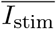 and duration *T*_*stim*_ is applied at time *T*_1_, *T*_2_, …, *T*_*N*_ (and otherwise zero), where *T*_*i*+1_ *> iT*_*stim*_ + *T*_*i*_. The values of 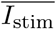 and *T*_*stim*_ are chosen to be those given by the default setting of the models. Similar to the voltage-clamp protocol parameterisation, equivalently we can define Δ*T*_*i*_ = *T*_*i*+1_ − *iT*_*stim*_ − *T*_*i*_, giving

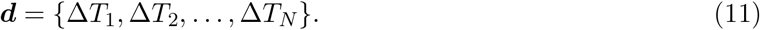

We further constrain Δ*T*_*i*_ ∈ [100, 2000] ms, ∀*i*, to make the designed experiments practically usable. We applied only the LSA designs for this experimental setting; to apply GSA designs for current clamp experiments, we would need to first explore the boundaries (not necessarily being a hypercube) of the parameter space that generate APs.

### 4.3 OED design measures

#### Local sensitivity analysis-based designs

The sensitivity matrix based on LSA, 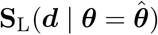,is defined as

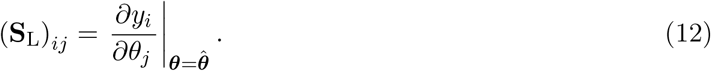

Here the subscript *i* (row) in *y*_*i*_ runs through all model outputs in ***y*** and all sampled time points *t*_*k*_^*^, and the subscript *j* (column) *θ*_*j*_ goes over all model parameters in ***θ***. The *Fisher information matrix* (FIM) is given by

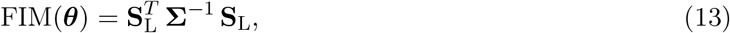

where **Σ** is the covariance matrix of the measurement data (noise), and we assume that data are a single output (a single current trace) with i.i.d. Gaussian noise through time such that **Σ** = *σ*^2^**I** with *σ* constant and **I** the identity matrix.

Many local design criteria are based on the parameter (co)variance matrix defined as

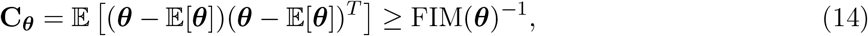

where 𝔼[·] denotes the expected value. This describes the covariance of the estimates of a fixed true parameter set^†^. For any unbiased estimator, where the expected value is equal to the true value of the parameter set, a lower bound for the variance is given by the inverse of the FIM (the inequality in Eq. (14)), known as the Cramér-Rao bound. Therefore, to evaluate the local design criteria based on the covariance matrix, we use the the Cramér-Rao bound and calculate FIM(***θ***)^−1^ instead (Walter and Pronzato, 1997; Schenkendorf et al., 2018). In fact, equality with the Cramér-Rao bound would be achieved as *K* (number of observations) approaches infinity

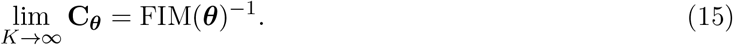

For proof of Eq. (15) see, for example, Pant (2018). Note, **C**_***θ***_, FIM, and **S**_*L*_ all depend on the choice of the system input ***d***.

A common class of the local design criteria is the ‘*alphabetic family*’ (Kiefer, 1959; Walter and Pronzato, 1997). These criteria each represent a cost function to be minimised, a subset of them (A-optimal, D-optimal, and E*-optimal designs, Atkinson and Donev, 1992) are given in Table 1, where *λ*_*max*_(**X**) and *λ*_*min*_(**X**), respectively, denote the maximum and minimum eigenvalues of a matrix **X**.

**Table 1:**
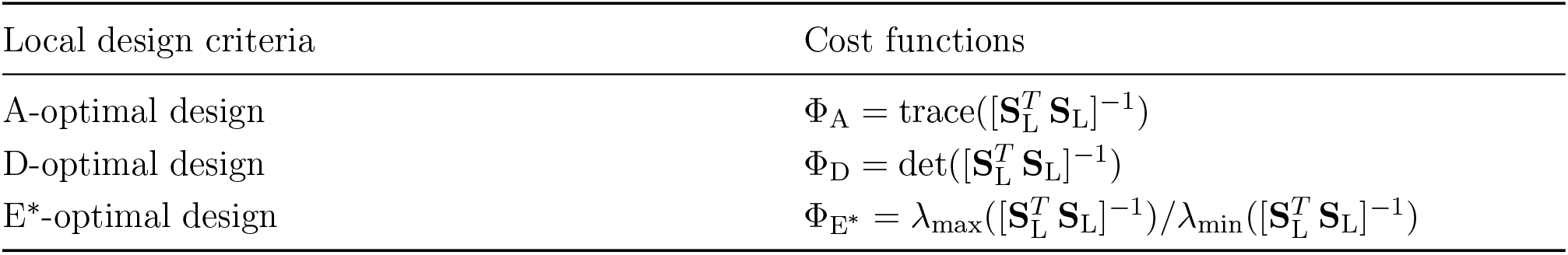
Local design criteria for OED. Here we have assumed the parameter covariance matrix **C**_***θ***_ for some given model parameters ***θ*** can be approximated as 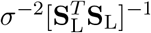, where *σ*^−2^ is dropped from the design criteria (cost functions) as it is assumed to be constant. E* is the modified E criterion.

| Local design criteria | Cost functions |
| --- | --- |
| A-optimal design | $\Phi_{\text{A}} = \text{trace}([\mathbf{S}_{\mathbf{L}}^T \mathbf{S}_{\mathbf{L}}]^{-1})$ |
| D-optimal design | $\Phi_{\text{D}} = \det([\mathbf{S}_{\mathbf{L}}^T \mathbf{S}_{\mathbf{L}}]^{-1})$ |
| E*-optimal design | $\Phi_{\text{E}^*} = \lambda_{\max}([\mathbf{S}_{\mathbf{L}}^T \mathbf{S}_{\mathbf{L}}]^{-1}) / \lambda_{\min}([\mathbf{S}_{\mathbf{L}}^T \mathbf{S}_{\mathbf{L}}]^{-1})$ |

The design criteria in Table 1 are parameter based criteria (Atkinson and Donev, 1992): these cost functions (as functions of the FIM) can be interpreted as properties (the shape and size) of the *confidence hyper-ellipsoid* (a generalisation of the confidence interval for multivariate statistics) for the parameters ***θ*** (Vanrolleghem and Dochain, 1998; Banga and Balsa-Canto, 2008). Therefore, by improving these properties of the confidence ellipsoid, the uncertainty of the inferred parameters using the data measured under these optimal design protocol would (in theory) be reduced.

One obvious limitation of the local designs is that they are *local*, i.e. they depend on a particular choice of the model parameters (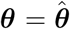in the equations above). Hence the design is only guaranteed to be optimal for that particular choice of parameters. However, we do not know the true model parameters in the experiments by definition. This issue could be alleviated by replacing the local sensitivity with an averaged sensitivity matrix

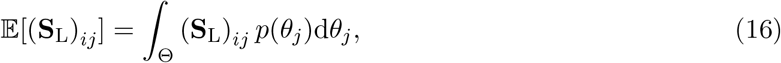

where *p*(*θ*_*j*_) is the probability density function of the *j*^*th*^ parameter, defined over a domain Θ (a subspace of the parameter space). The integral in Eq. (16) can be approximated using Monte Carlo simulations. Although this averaged sensitivity matrix has been shown to give a more robust design (Chu and Hahn, 2013), it is does not take the multivariate interaction into account (Schenkendorf et al., 2018), and hence the framework of GSA comes into play.

#### Global sensitivity analysis-based designs

One of the most straight forward ways of doing GSA-based design is by replacing the **S**_*L*_ in Table 1 with a sensitivity matrix based on GSA methods (Rodriguez-Fernandez et al., 2007; Kucerová et al., 2016; Schenkendorf et al., 2018). We can use variance-based methods such as the *Sobol method* (also known as the *Sobol indices* Sobol, 2001). The Sobol method decomposes the variance of the model output into fractions, and attributes these variances to parameters or sets of parameters, usually referred to as the *order*. The first-order Sobol index attributes these variances to each parameter *θ*_*j*_ alone; whilst higher order Sobol indices attribute the variances to different combinations of parameters. These variances are calculated using a set of parameters sampled within a domain Θ, a subspace of the parameter space. These parameters can be sampled using various methods, for example Monte Carlo methods, or some low-discrepancy sequences (including Sobol sequences or Latin hypercube sampling); we used the extension of the Sobol sequence by Saltelli (2002) and Saltelli et al. (2010) here. The first-order Sobol sensitivity index is given by

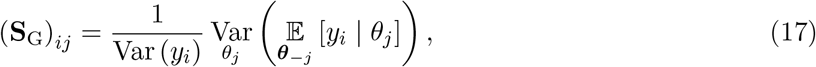

where Var (*y*_*i*_) is the total variance of the model output 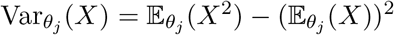 denotes the conditional variance only over parameter *θ*_*j*_, and 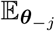 indicates the expected value taken over all parameters ***θ*** except parameter *θ*_*j*_. Criteria using **S**_*G*_ are given in Table 2. Note that one may further include higher order Sobol indices for analysing multivariate parameter interactions and dependencies (Kucerová et al., 2016), but since we would like to maximise the independent sensitivity of the parameters for the experimental design, using the first-order sensitivities is sufficient.

**Table 2:** Global design criteria for OED based on a generalisation of the local design criteria in Table 1 using a GSA method.

| GSA-based design criteria | Cost functions |
| --- | --- |
| A-optimal design | $\Phi_A^{\text{GSA}} = \text{trace}([\mathbf{S}_G^T \mathbf{S}_G]^{-1})$ |
| D-optimal design | $\Phi_D^{\text{GSA}} = \det([\mathbf{S}_G^T \mathbf{S}_G]^{-1})$ |
| E*-optimal design | $\Phi_{E^*}^{\text{GSA}} = \lambda_{\max}([\mathbf{S}_G^T \mathbf{S}_G]^{-1}) / \lambda_{\min}([\mathbf{S}_G^T \mathbf{S}_G]^{-1})$ |

### 4.4 Optimisation of the protocols

To perform the optimisation of the voltage-clamp protocol as defined by Eq. (3), we optimise the design measures (cost functions) by varying the protocol parameters ***d*** in Eq. (10) for voltage clamp and in Eq. (11) for current clamp. The protocol was initialised with parameters ***d*** = ***d***_0_ randomly sampled uniformly within the defined boundaries (the protocol parameter space). Optimisations were performed using the covariance matrix adaptation evolution strategy (Hansen, 2006) via our open source Python package, PINTS (Clerx et al., 2019b). This was repeated 10 times with different initial protocol parameters, and the best result out of all repeats was used and presented in the results.

To compute the Sobol indices, we define a hypercube for the model parameter space Θ, where each conductance value *g*_*j*_ in Eq. (6) can be scaled from *e*^−2^ ≈ 0.14 to *e*^2^ ≈ 7.39 times its original value. We use the Saltelli et al. (2010) extension of a Sobol sequence to generate 512 parameter samples within this parameter space Θ for computing the Sobol indices.

To numerically calculate the local sensitivity matrix **S**_*L*_, we use the first-order central difference to approximate the local derivatives with a step-size of 0.1% of the parameter values. It may be worth noting that although LSA-based criteria require local derivatives, we are not truly interested in (local) infinitesimal changes when designing the protocols, hence a reasonable approximation of the derivatives is typically enough.

### 4.5 Comparing the performance of OEDs

#### Cross-measure evaluations

We defined a cross-measure matrix **X** for each optimised protocol, such that each entry *X*_*i,j*_ was the criterion *j* score for this protocol with currents generated by model *i*, with 1 *< j < N*_*measure*_ and

1 *< i < N*_*model*_. The models indexed by *i* were the ten Tusscher et al. (2004) model, the Fink et al. (2008) model, the O’Hara et al. (2011) model, the Chang et al. (2017) model, the Tomek et al. (2019) model, and the averaged model; the O’Hara et al. (2011) model and the averaged model were used to optimise the protocols, and the remaining five were included to assess how these optimised protocols perform for experiments from different models. The criteria indexed by *j* were the LSA A-, D-, E*-designs, and the GSA A-, D-, E*-designs, respectively, for the voltage-clamp mode; *j* were the LSA A-, D-, E*-designs, respectively, for the current-clamp mode.

The numerical values for different cost functions were not directly comparable, so we normalized the values. Normalisation should be better than using a simple ranking as it should highlight outliers better. Outliers concerned us here, because we expected the best protocols to be robust to the model (possible current kinetics in reality) and to perform well under all measures. Ideally, we would like to have a protocol that is good under all criteria and all possible models of kinetics. The normalisation for each entry was calculated using 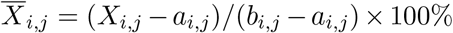, where *a*_*i,j*_ and *b*_*i,j*_ were the minimum (best) and maximum (worst) scores seen across all the protocols (Protocols A–L for voltage clamp and Protocols O–T for current clamp). Note the normalisation (percentile) was calculated with only the optimised protocols, i.e. without the benchmark protocols, for clarity. Therefore the percentiles shown in Figures 6 and 7 were not necessarily within 0 to 100%. at the entry *i, j* (i.e. with this criterion and current model).

#### Practical evaluations

We further evaluated the optimised protocols from a practical point of view. We generated synthetic data under these protocols with synthetic noise, and we assessed identifiability of the parameters in Eq. (6).

We generated synthetic data using the O’Hara et al. (2011) model with ∼𝒩 (0, *σ*^2^) as the synthetic noise, where *σ* = 0.15 A F^−1^ for voltage clamp and *σ* = 0.15 mV for current clamp. Then the same model was used to fit the synthetic data, to test the ideal case where we know the ground truth kinetics. We reparameterised the models with a scaling factor *s*_*j*_ for the maximum conductance *g*_*j*_, where *s*_*j*_ = 1 was the original literature value. The likelihood of observing the data **Y** with *K* time points given the model parameters ***θ*** was

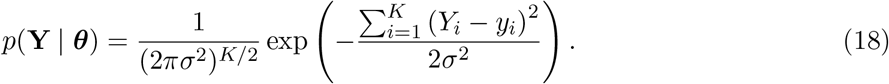

The posterior distribution of the parameters was

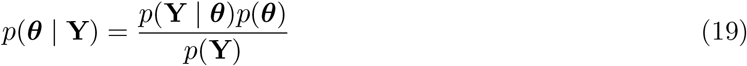

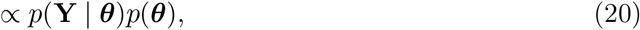

where *p*(***θ***) is the prior, and *p*(**Y**) is the marginal likelihood which is a constant. Uniform priors were used 𝒰 (0.04, 25) for the scaling of the conductance values, which was wider than the subspace we used for computing the GSA to test its robustness for parameters outside the design parameter space.

The posterior distributions of the parameters were estimated using a Monte-Carlo based sampling scheme—a population MCMC (Jasra et al., 2007) algorithm with adaptive Metropolis as the base sampler—via our open source Python package, PINTS (Clerx et al., 2019b). We ran four chains for the population MCMC, each with 4 × 10^4^ samples, and discarded the first 10^4^ samples as warm-up period.

## Acknowledgments

This work was performed in part at the High Performance Computing Cluster supported by the Information and Communication Technology Office of the University of Macau.

## Author Contributions

Conceptualization: CLL, MC, DJG, GRM. Methodology: CLL, GRM. Investigation: CLL. Writing— original draft: CLL, GRM. Writing—review: CLL, MC, DJG, GRM.

## S1 Supplementary materials

**Figure S1:**
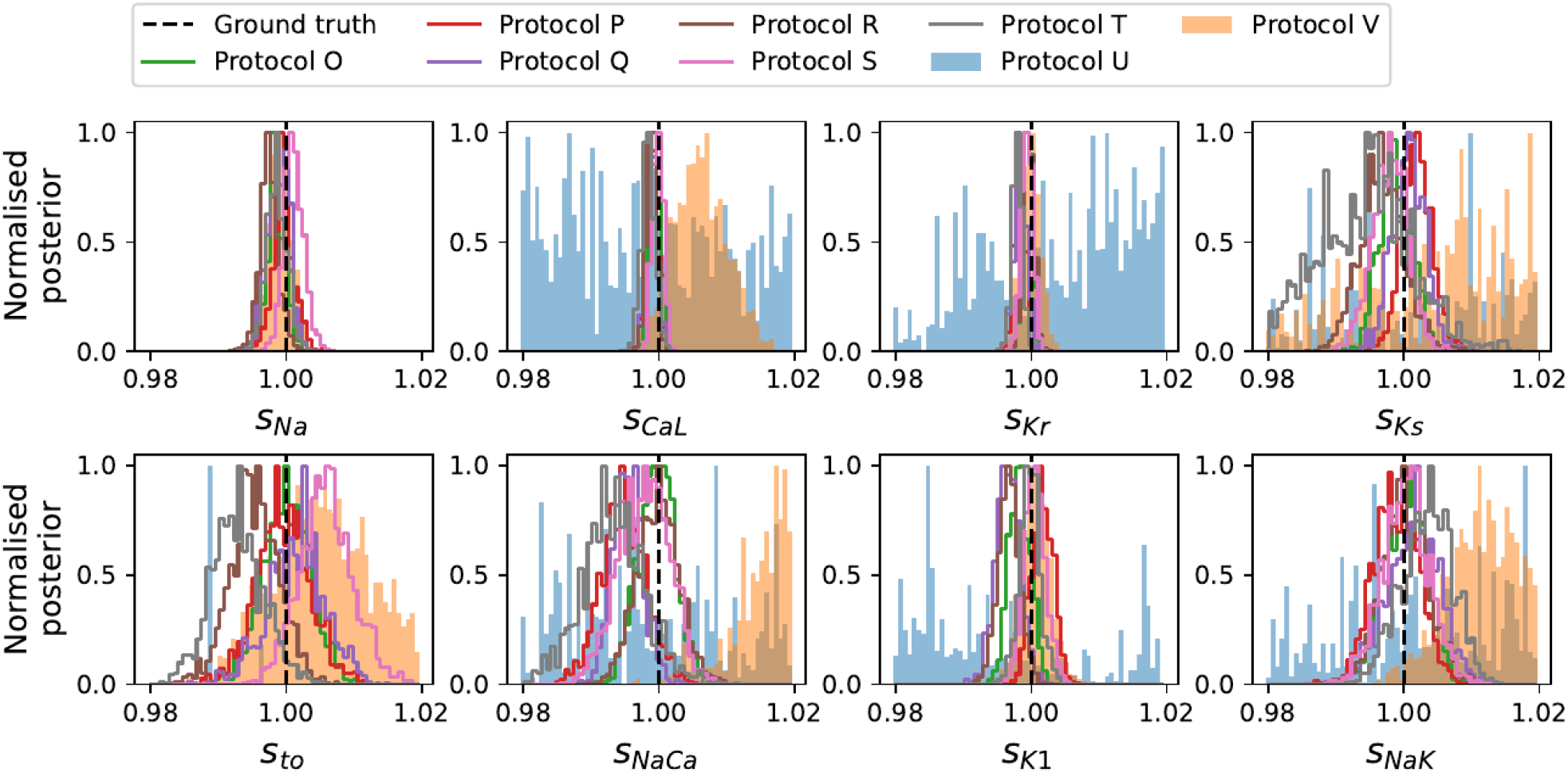
The marginal posterior distribution of the parameters for the synthetic study with current clamp, generated with the O’Hara et al. (2011) model and fitted with the same model. A magnification version of Figure 5.

**Figure S2:**
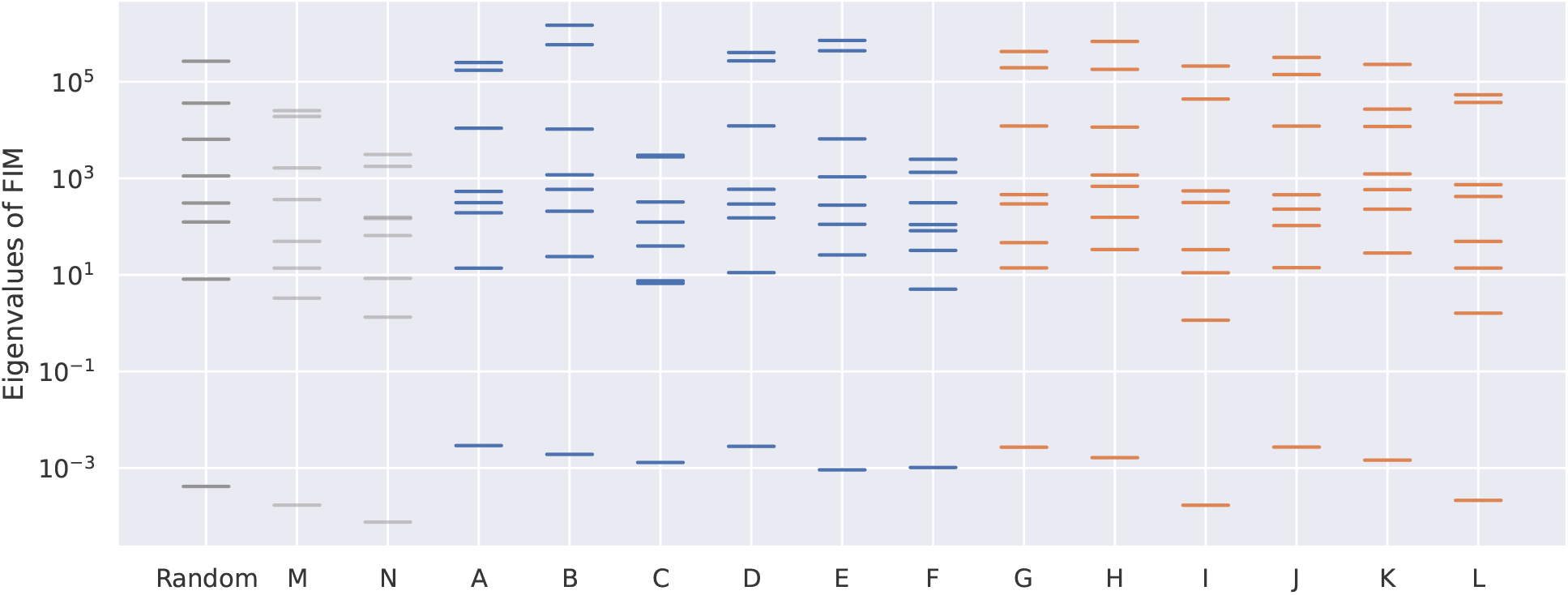
The eigenvalue spectra of the Fisher information matrix (FIM) for the voltage clamp protocols presented in the main text.

## Footnotes

* That is, for *N*_*o*_ model outputs and *N*_*t*_ time points, *i* = 1, 2, …, *N*_*o*_, *N*_*o*_ + 1, *N*_*o*_ + 2, …, *N*_*o*_ *× N*_*t*_.

† Note that under this framework, the true parameters do not vary, as opposed to the Bayesian framework where the true parameters can be treated as random variables.

